## Supplemental figures for "Spatial transcriptomics and single-nucleus RNA sequencing reveal a transcriptomic atlas of adult human spinal cord"

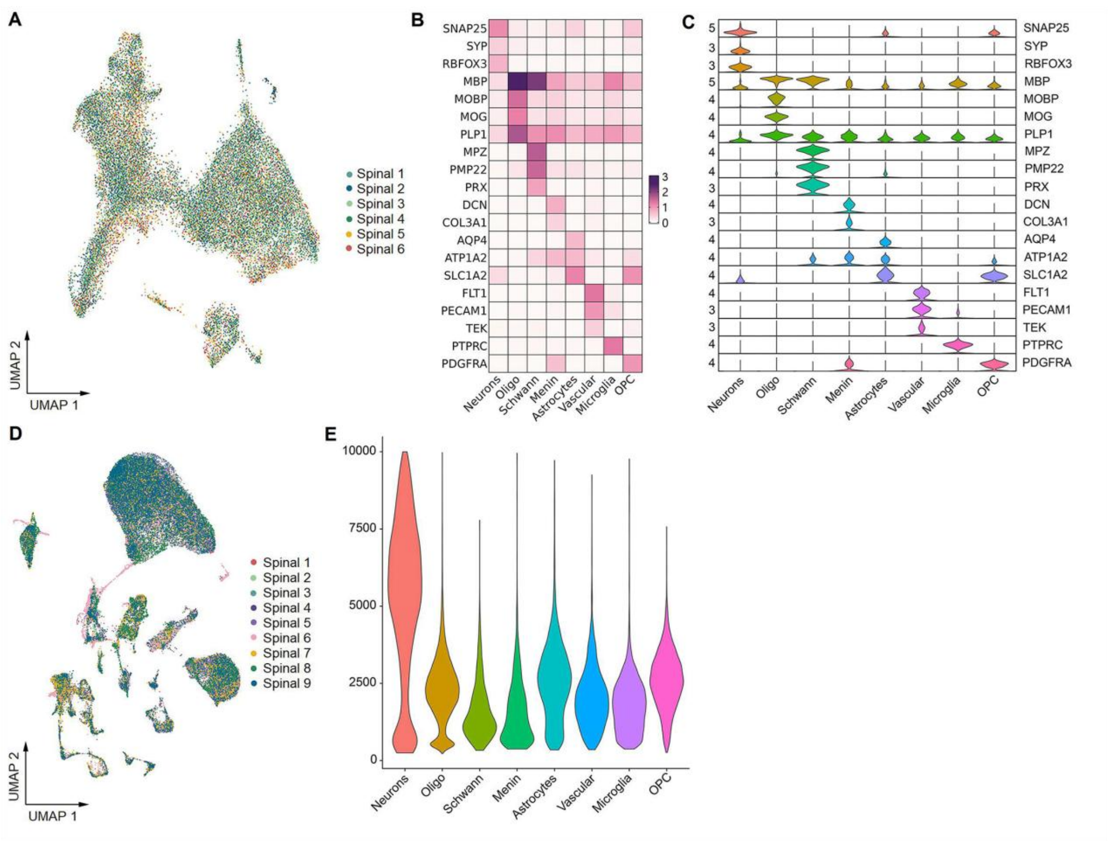


**Supplemental figure 1.** (**A**) UMAP plot showing the contribution of each donor to spinal cluster formation by spatial transcriptomics. (**B-C**) Heatmap (B) and violin plot (C) showing the expression of selected marker genes across all eight cell types in the spinal cord by single-nucleus RNA-seq. (**D**) UMAP plot showing the contribution of each donor to spinal cluster formation by single-nucleus RNA-seq. (**E**) Violin plot showing genes detected per nucleus in each spinal cell type by single nucleus RNA-seq. Oligo, oligodendrocytes; OPC, oligodendrocyte precursor cells; Menin, meningeal cells; UMAP, uniform manifold approximation and projection.

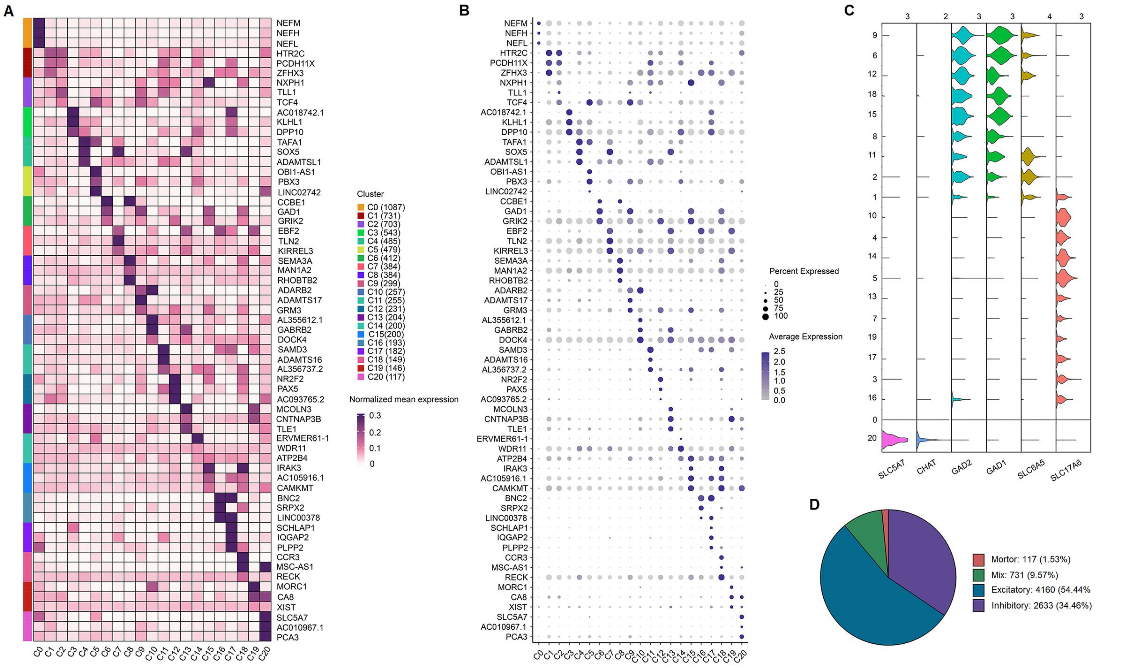


**Supplemental figure 2.** (**A**) Heatmap showing the expression of the top 3 most differentially expressed genes across different human spinal neuronal clusters. (**B**) Dot plot showing the expression of the top 3 most differentially expressed genes across different spinal neuronal clusters. (**C**) Violin plot showing the expression of excitatory, inhibitory, and cholinergic marker genes across all 21 neuronal clusters in the human spinal cord. (**D**) The component percentage of different clusters according to neurotransmitter status. Mix, mixed excitatory and inhibitory clusters.

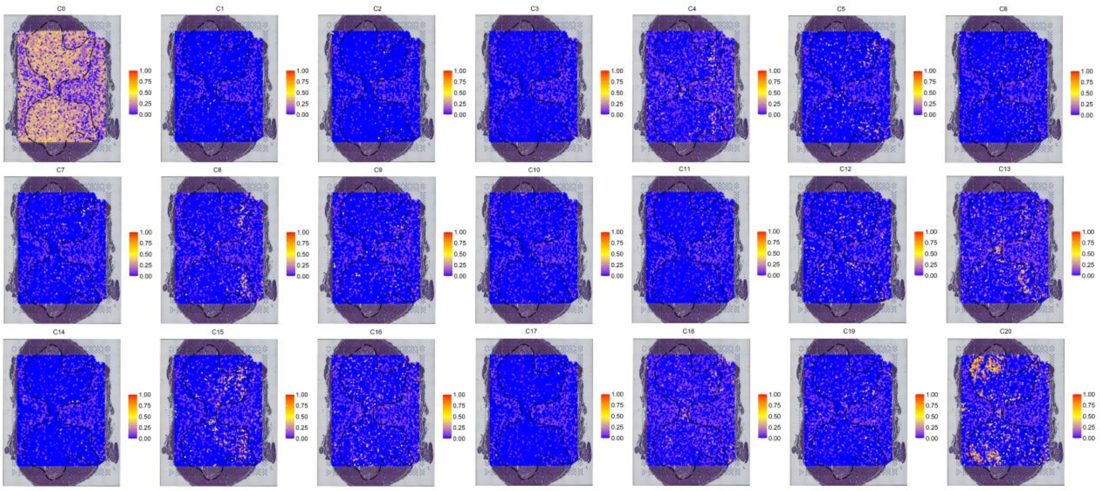


**Supplemental figure 3.** The spatial spot showing the distribution pattern of 21 neuronal clusters in the human spinal cord. The color scale represents the probable percentage of single nucleus RNA-seq data that mapped to spatial spots. For a specific spot, the sum of the probable percentage of C0-C20 was defined as 1.

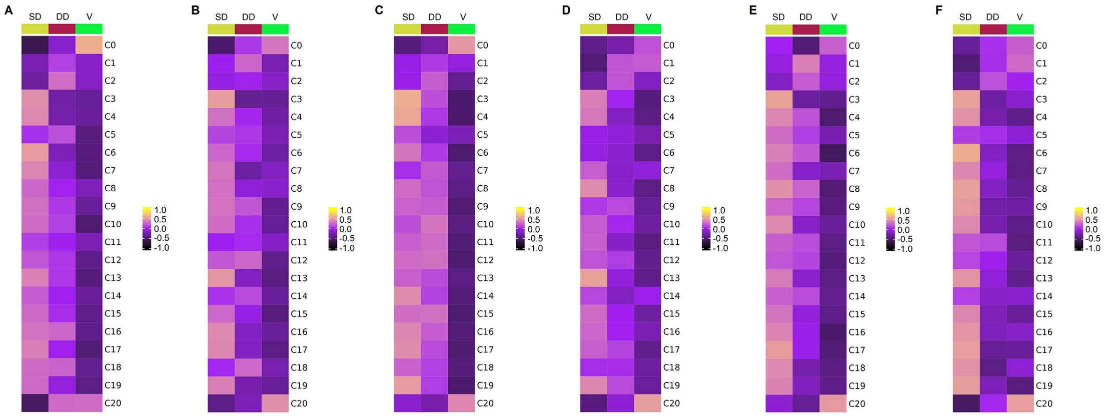


**Supplemental figure 4. (A-F)** Gene set variation analysis (GSVA) showing the spatial distribution patterns of neuronal clusters in different subregions of coronal sections from the lumber spinal cord of each human donor. The color scale represents the enrichment scores. SD, superficial dorsal horn; DD, deep dorsal horn; V, ventral horn.

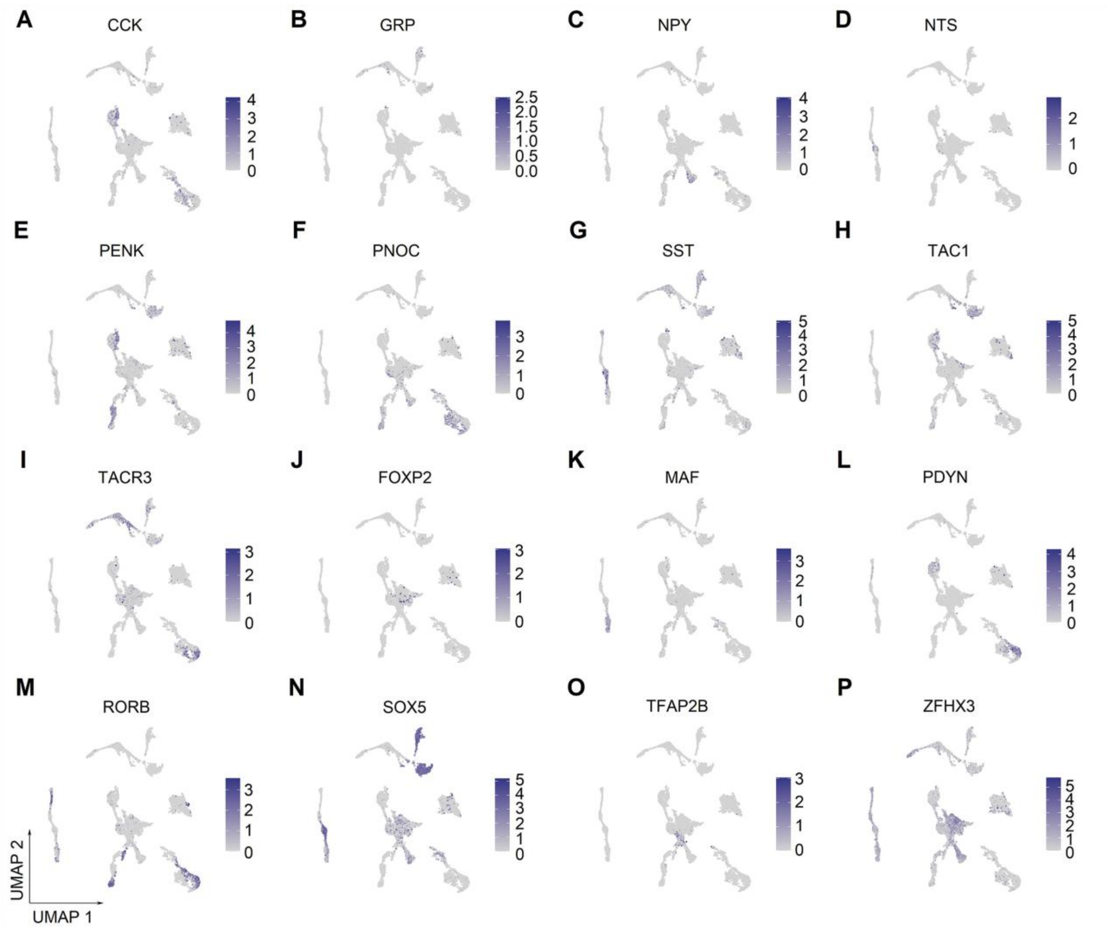


**Supplementalal figure 5.** UMAP plot showing the expression of representative marker genes in human spinal neuronal clusters. UMAP, uniform manifold approximation and projection.

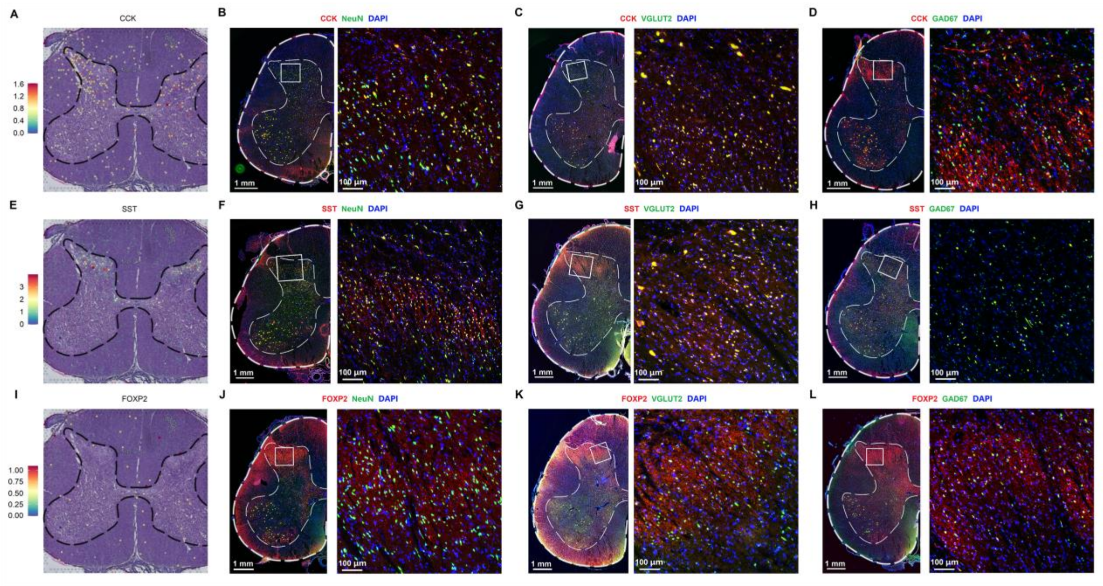


**Supplementalal figure 6.** (**A**) Representative section showing the spatial distribution of CCK in the human spinal cord. (**B**) Representative immunofluorescence images of CCK in human spinal neurons. (**C-D**) Representative immunofluorescence images of CCK in excitatory (VGLUT2, C) and inhibitory (GAD67, D) neurons of the human spinal cord. (**E**) Representative section showing the spatial distribution of SST in the human spinal cord. (**F**) Representative immunofluorescence images of SST in human spinal neurons. (**G-H**) Representative immunofluorescence images of SST in excitatory (VGLUT2, G) and inhibitory (GAD67, H) neurons of the human spinal cord. (**I**) Representative section showing the spatial distribution of FOXP2 in the human spinal cord. (**J**) Representative immunofluorescence images of FOXP2 in human spinal neurons. (**K-L**) Representative immunofluorescence images of FOXP2 in excitatory (VGLUT2, K) and inhibitory (GAD67, L) neurons of the human spinal cord.

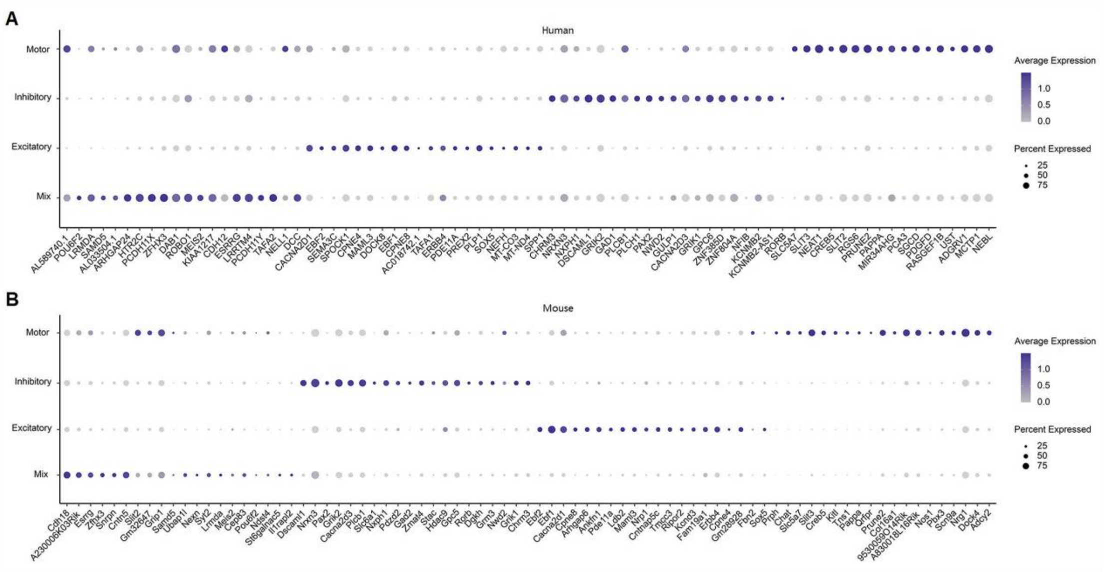


**Supplementalal figure 7.** (**A-B**) Dot plot showing the expression of the top 20 most differentially expressed genes between excitatory, inhibitory, and cholinergic clusters of the spinal cord in humans (A) and mice (B).

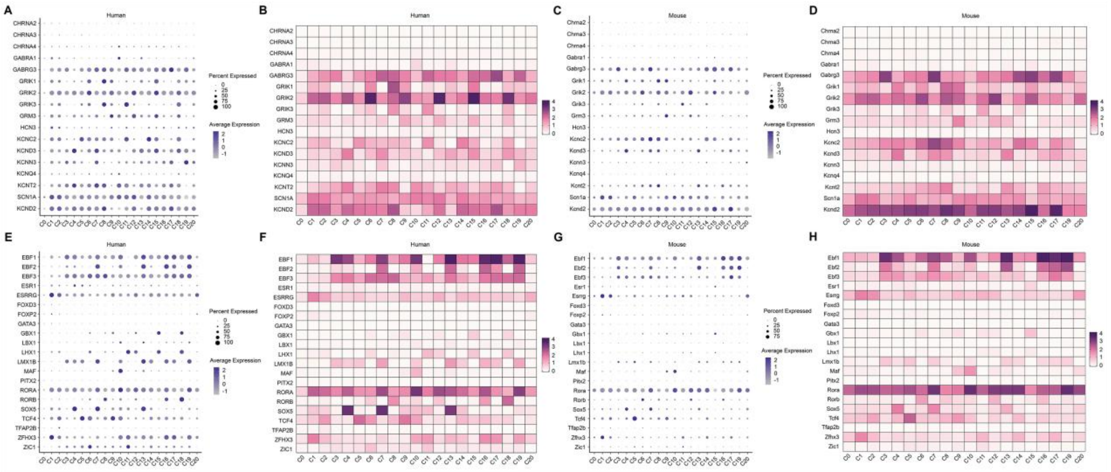


**Supplementalal figure 8.** (**A-B**) Dot plot (A) and heatmap (B) showing the expression of classic ion channels in human spinal neuronal clusters. (**C-D**) Dot plot (C) and heatmap (D) showing the expression of classic ion channels in mouse spinal neuronal clusters. (**E-F**) Dot plot (E) and heatmap (F) showing the expression of classic transcription factors in human spinal neuronal clusters. (**G-H**) Dot plot (G) and heatmap (H) showing the expression of classic transcription factors in mouse spinal neuronal clusters.

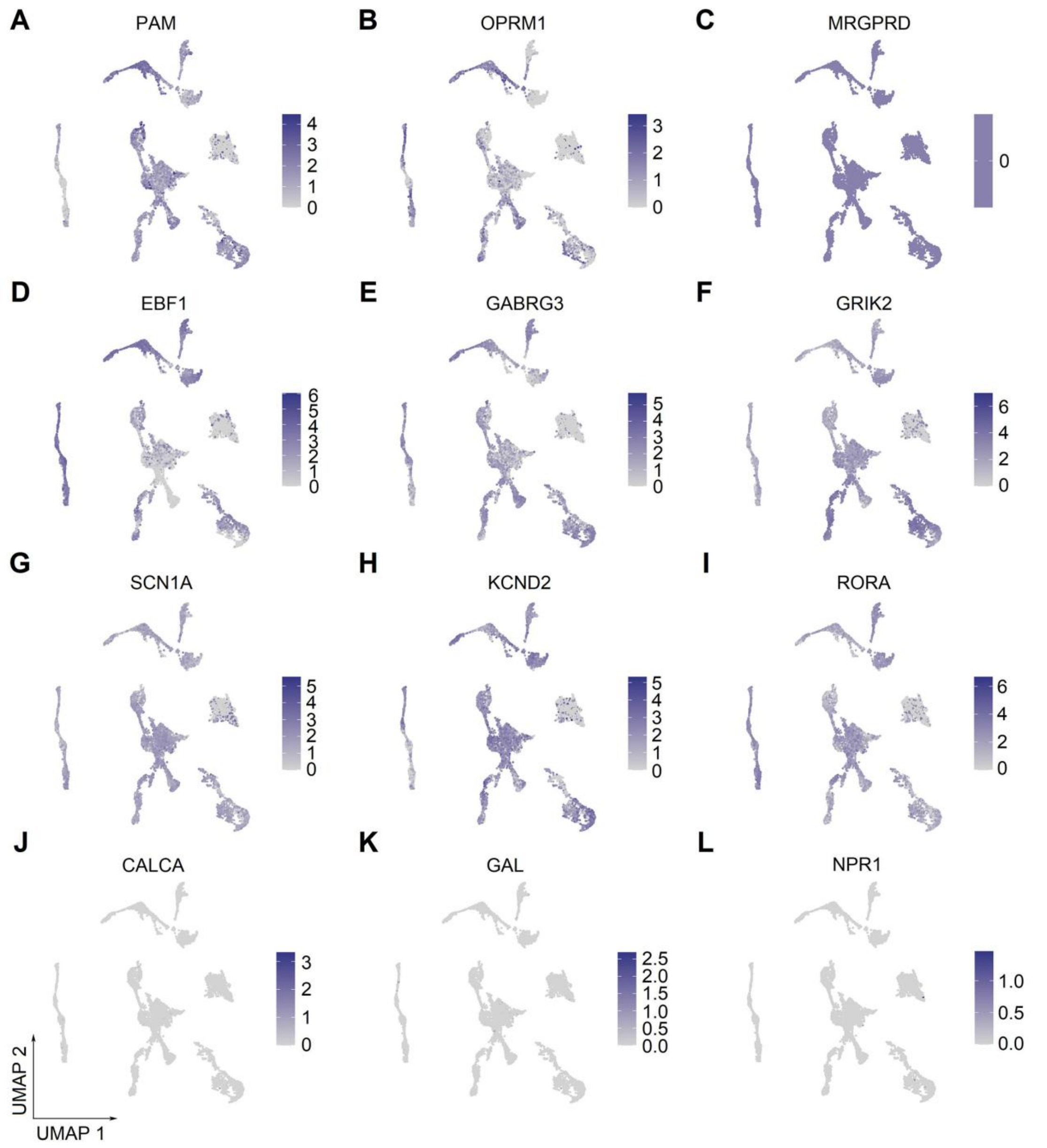


**Supplementalal figure 9.** (**A-L**) UMAP plot showing the broad (A-I) or low (J-L) expression of selected marker genes in human spinal neuronal clusters. UMAP, uniform manifold approximation and projection.

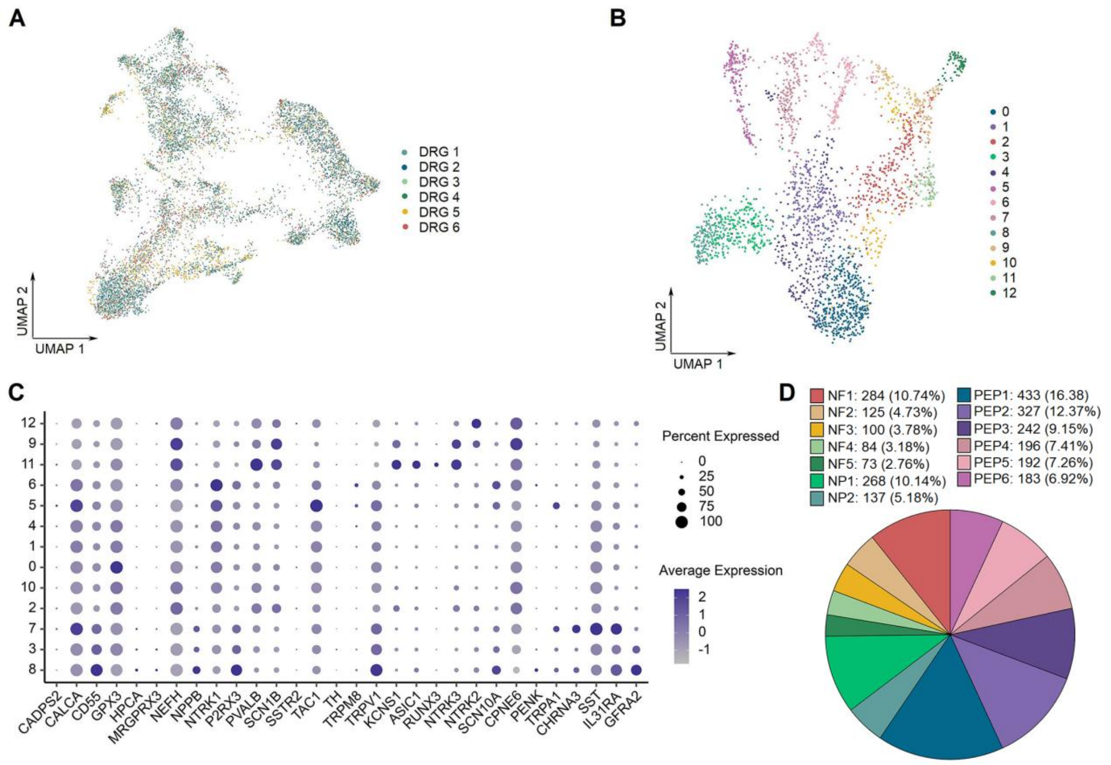


**Supplementalal figure 10.** (**A**) UMAP plot showing the contribution of each donor to cluster formation in the DRG by spatial transcriptomics. (**B**) UMAP plot showing 13 types of human DRG neurons. Dots, individual spots; colors, cell types. (**C**) Dot plot showing the expression of representative marker genes across all DRG neuronal subtypes. (**D**) The component percentage of DRG neuronal subtypes. UMAP, uniform manifold approximation and projection.

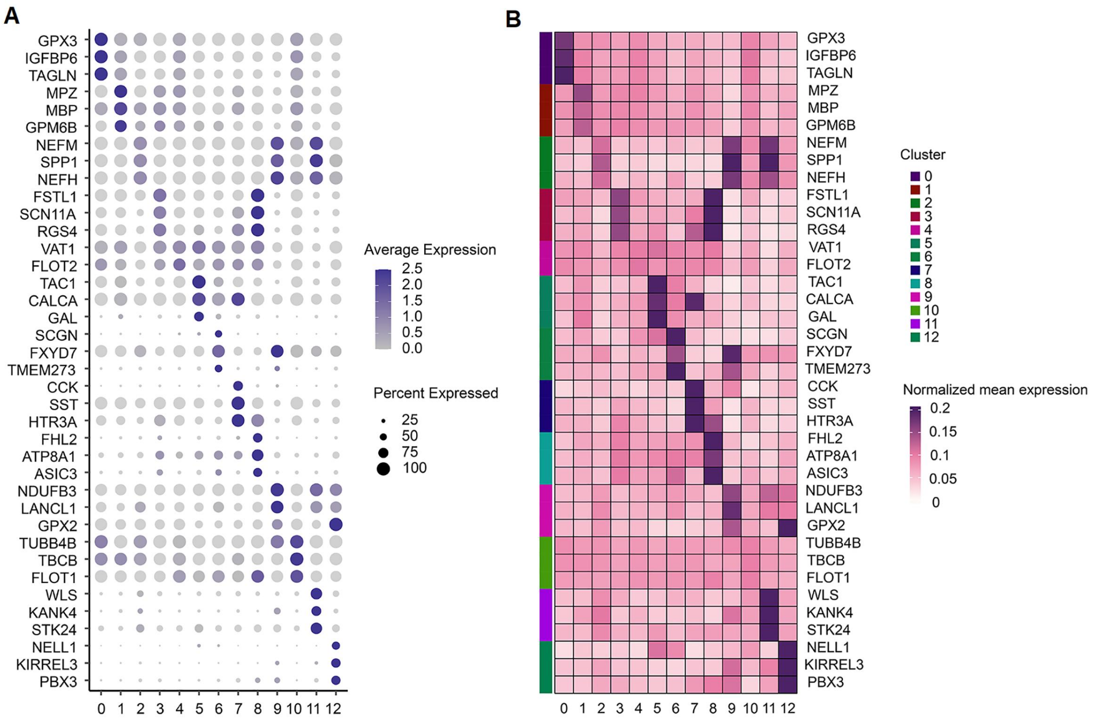


**Supplementalal figure 11.** (**A-B**) Dot plot (A) and heatmap (B) showing the expression of the top 3 most differentially expressed genes across human DRG neuronal subclusters.

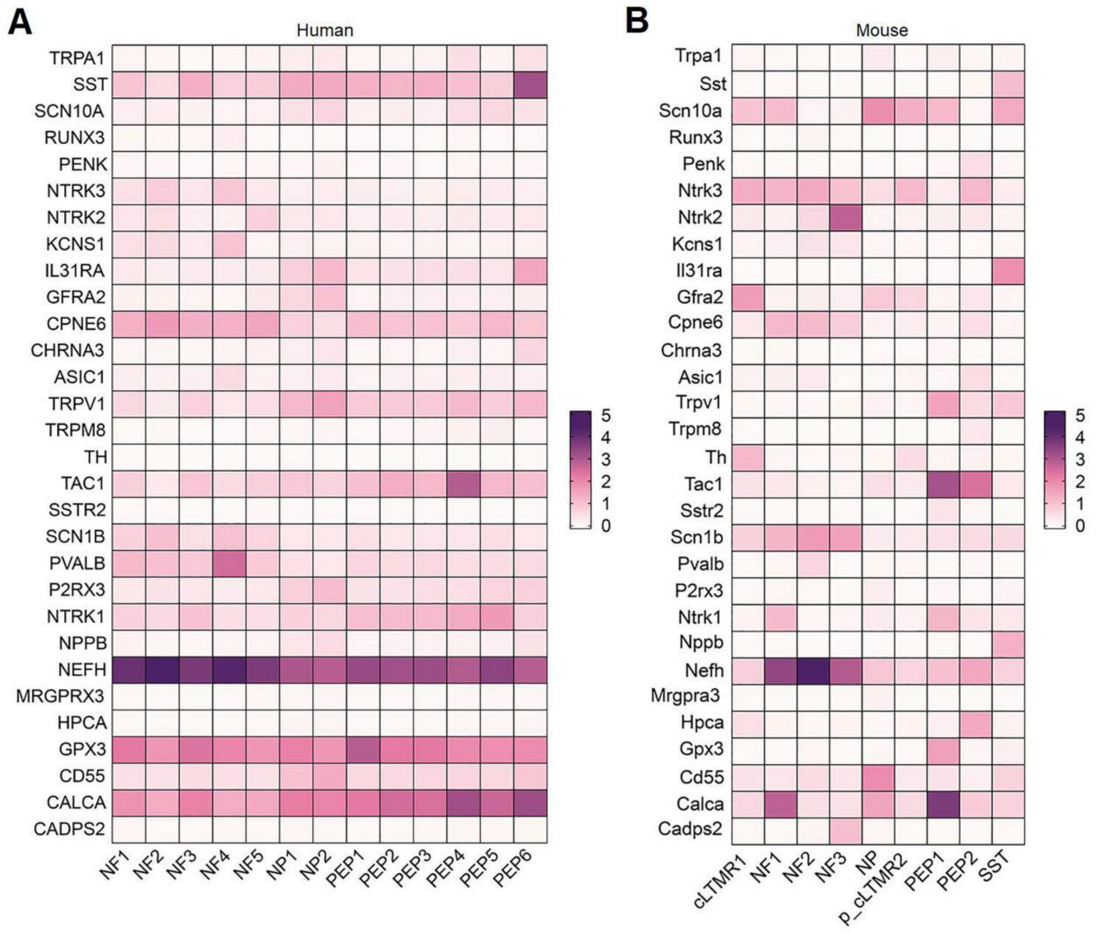


**Supplementalal figure 12.** (**A-B**) Heatmap showing the expression of classical marker genes in human (A) and mouse (B) DRG neuronal clusters.

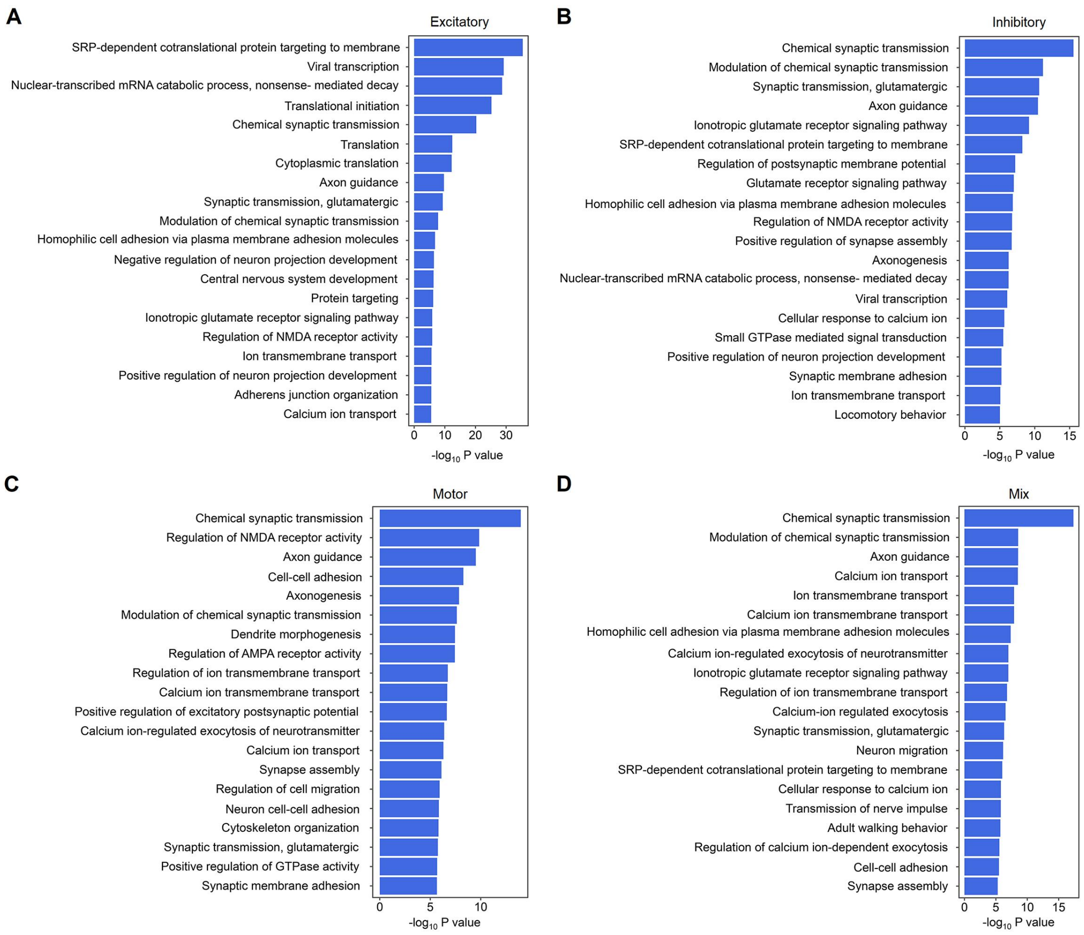


**Supplementalary figure 13.** (**A-D**) Summarized GO terms among the top genes in excitatory (A), inhibitory (B), motor (C), and mixed excitatory and inhibitory (D) clusters of the human spinal cord. GO, Gene Ontology;
